# Is hybridization evil? –– Competition generates alternative stable states of coexistence of hybridizing species

**DOI:** 10.1101/2025.11.26.690876

**Authors:** Keiichi Morita, Ryo Yamaguchi

**Affiliations:** Graduate School of Environmental Science, Hokkaido University, Sapporo, Hokkaido 060-0810, Japan; Interdisciplinary Theoretical and Mathematical Sciences Program, Institute of Physical and Chemical Research, RIKEN, 2-1 Hirosawa, Wako, Saitama 351-0198 Japan; Department of Evolutionary Studies of Biosystems, School of Advanced Sciences, The Graduate University for Advanced Studies, SOKENDAI, Hayama, Kanagawa 2400193, Japan; Department of Advanced Transdisciplinary Science, Hokkaido University, Sapporo, Hokkaido 060-0808, Japan; Department of Zoology & Biodiversity Research Centre University of British Columbia Vancouver British Columbia Canada

**Keywords:** hybridization-mediated coexistence, alternative stable states, backcrossing

## Abstract

Predicting ecological outcomes of hybridizing species remains challenging because of complex interactions between genetic and ecological processes. Previous studies have mainly focused on exclusion scenarios driven by genetic mechanisms, whereas few have explored the impact of competitive interactions among parent species and their hybrids on generating multiple coexistence regimes. Here, we develop a population dynamics model that explicitly couples hybridization genetics with population dynamics driven by competitive interactions among two parent species and their hybrids. Our analysis reveals that backcrossing can fundamentally reshape coexistence regimes, generating five alternative outcomes: (i) alternative stable states of coexistence of both parent species and hybrids, (ii) frequency-independent coexistence, (iii) frequency-dependent coexistence, (iv) competitive exclusion of a rarer species, and (v) extinction of all populations. These outcomes arise from the balance between two key processes: regeneration of parental genotypes through hybrid × hybrid mating and competitive exclusion mediated by hybrids. Thus, hybridization can generate multiple stable coexistence regimes even when species would otherwise be excluded. This eco-genetic mechanism provides a unifying theoretical basis for predicting the ecological consequences of hybridization, highlighting the crucial role of competitive interactions in determining coexistence outcomes.

## 1. Introduction

Understanding the mechanisms that maintain coexistence of closely related species remains a central challenge in community ecology. Classical theories predict that rarer species are likely excluded through competitive exclusion driven by niche overlap (Chesson 2000, Levine and HilleRisLambers 2009, Mittelbach and McGill 2019). However, empirical studies frequently reveal diverse coexistence patterns in hybrid zones, posing a challenge to traditional competition theories (Arntzen et al. 2017, Mandeville et al. 2019, 2022, Martins et al. 2024).

Hybridization––the interbreeding of genetically distinct, closely related populations––is increasingly recognized as a novel ecological process shaping coexistence and community structure (Mallet 2007, Abbott et al. 2013, Todesco et al. 2016, Porretta and Canestrelli 2023). Accumulating empirical evidence demonstrates that hybridization between closely related species is widespread, with backcrossing to parent species occurring more frequently than previously appreciated (Rieseberg et al. 2003, Taylor and Larson 2019, Edelman et al. 2019). This prevalence has increased in the Anthropocene, as climate-driven range shifts, habitat modification, and human-assisted introductions have created novel opportunities for secondary contact between previously isolated populations (Rhymer and Simberloff 1996, Ottenburghs 2021). Consequently, understanding the ecological consequences of hybridization has become critical for predicting community dynamics in changing environments.

Hybridization often causes the extinction of rare species, raising urgent concerns in conservation management (Levin et al. 1996, Rhymer and Simberloff 1996), yet it also shapes biological invasion, coexistence of two closely related species, hybrid-zone movement and shifts in species distributions (Barton and Hewitt 1989, Buggs 2007, Schierenbeck and Ellstrand 2009, Pfennig et al. 2016, Yamaguchi et al. 2019, Irwin and Schluter 2022). Despite this broad influence, predictive theory linking hybridization to community-level outcomes remains underdeveloped, largely because ecological and genetic processes interact in complex ways.

Community ecologists have considered that hybridizing species interact through both reproductive and ecological pathways. In cases where interspecific mating produces inviable or sterile offspring, wasted reproductive effort reduces per capita growth rates of both species (i.e., reproductive interference or demographic swamping) (Kuno 1992, Kishi and Nakazawa 2013, Todesco et al. 2016, Kyogoku 2020). Reproductive interference drives sexual exclusion of rarer species due to positive frequency-dependence in an ecological community (Gröning and Hochkirch 2008, Kyogoku 2020). Moreover, resource competition may elevate the risk of competitive exclusion beyond scenarios involving reproductive interference alone (Kuno 1992, Kishi and Nakazawa 2013, Kyogoku 2020). When hybrids are fertile, however, ecological dynamics become far more complex and likely unpredictable, as hybrids interact competitively and reproductively with parent species. Empirical research has highlighted the importance of competitive interactions in hybrid zones (Vallin et al. 2012), and has observed competition between parent species and hybrids across various taxa (Boersma 1995, Ayres et al. 2004, Seiler and Keeley 2007). Despite these findings, the development of extended reproductive-interference models that incorporate competitive interactions among parent species and hybrids has not kept pace.

Hybridization genetics also plays a substantial role in determining coexistence outcomes, particularly for rare species. When hybrid fitness reduction is mild, hybridization can lead either to genetic swamping—replacement of resident genomes by introgressed alleles—or to genetic rescue of endangered populations (Epifanio and Philipp 2000, Prentis et al. 2007, Donovan et al. 2010, Todesco et al. 2016, Williams et al. 2019, Ålund et al. 2024). Thus, hybridization has the dual potential to both promote and prevent coexistence.

Despite advances in understanding both ecological and genetic mechanisms, theoretical frameworks that unify these mechanisms to predict when hybridization promotes or prevents coexistence remain underdeveloped. Most theoretical studies have focused on genetic processes leading to extinction (Huxel 1999, Epifanio and Philipp 2000, Wolf et al. 2001, Ferdy and Austerlitz 2002, Hall and Ayres 2009). Previous models of reinforcement and reproductive character displacement incorporate population dynamics but generally assume coexistence, rather than identifying the conditions under which extinction occurs (Liou and Price 1994, Goldberg and Lande 2006, Konuma and Chiba 2007, Kyogoku and Yamaguchi 2023). Although several recent studies incorporating competitive interactions among hybrids and parent species have documented frequency-dependent coexistence and multiple stable states (Quilodrán et al. 2018, Irwin and Schluter 2022, Reed et al. 2024), the conditions that generate these outcomes in parent–species–hybrid systems remain poorly understood. In particular, the conditions under which hybridization facilitates coexistence—either through genetic rescue or through the balance of hybridization genetics and parent–hybrid competition—remain largely unexamined (see Wolf et al. (2001) for a related discussion of hybrid competitive ability affecting resident-species persistence).

In this study, we address this gap by developing a population dynamics model that explicitly couples the population genetics of hybridization with competitive interactions between all possible pairings—between parent species, between parents and hybrids, and among hybrids. By analyzing scenarios with and without viable hybrids, we identify the ecological conditions under which backcrossing facilitates coexistence, in contrast to previous studies that have focused primarily on exclusion outcomes. For instance, Irwin and Schluter (2022) categorized ecological outcomes of hybridization into five types, one being coexistence of two closely related species, and the other four involving extinction of at least one species, whereas our study identifies three distinct types of coexistence. These findings provide a mechanistic basis for understanding coexistence patterns in natural hybrid systems and offer testable predictions for empirical studies.

## 2. Model

We investigate ecological outcomes when a closely related species colonizes a resident species’ habitat, focusing on conditions enabling species coexistence versus exclusion on secondary contact. We consider the ecological dynamics of two hybridizing species and their hybrid population prior to the completion of reinforcement or reproductive character displacement. Our mathematical framework extends the Lotka-Volterra competition model by explicitly incorporating population genetics of assortative mating, reproductive interference, and hybridization. In doing so, we construct an updated version of reproductive interference models that incorporates population dynamics of a hybrid population (Kuno 1992, Kishi and Nakazawa 2013, Schreiber et al. 2019). We first analyze the community dynamics without backcrossing (Result 3.1), then analyze conditions for competitive exclusion where a rarer species and a hybrid population go extinct (Result 3.2.1) and various coexistence scenarios (Result 3.2.2–3.2.4).

We explore one-locus model to focus on the role of competitive interactions in coexistence. We consider that a single locus pleiotropically controls both mating cues (e.g., male ornamentation) and the corresponding species recognition traits. Such traits are often referred to as a “self-referencing traits” (Hauber and Sherman 2001, Kopp et al. 2018). To isolate the combined effect of hybridization genetics of self-referencing traits and competitive interactions, we do not include ecological traits related to resource competition and other genetic mechanisms affecting such traits. Throughout, our model assumes two diploid species, overlapping generations, a single reproductive event per female per lifetime, and a 1:1 offspring sex ratio.

The resident species (species 1) is fixed for genotype AA, whereas invasive species (species 2) is fixed for genotype aa. Hybrid offspring, resulting from interspecific mating, have the genotype Aa. The mating trait value is determined by the single locus with two alleles. The effects are additive: allele A contributes a value of 1 to the trait, and allele a contributes a value of 0. As a result, the trait value for species 1, 𝑥_1_, species 2, 𝑥_2_, and hybrid, 𝑥_h_ are 2, 0, and 1, respectively. Mating probability between a female of species *i* and a male of type *j* is determined by the difference in their mating trait values, following a Gaussian function:

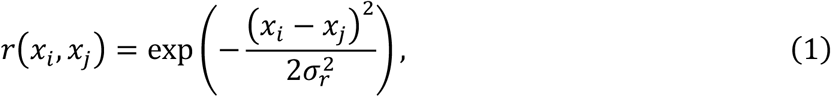

where 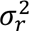 represents the inverse of the strength of female mate choice, choosiness or conspecific mate preference (we call female ‘choosiness’ hereafter) (Doebeli 2005, Kopp et al. 2018, Irwin and Schluter 2022). A smaller 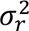 indicates stronger (i.e., more precise) assortative mating preference.

Model (i): Reproductive interference model

The model is described by a system of Ordinary Differential Equations (ODEs). Let *N*_1_, *N*_2_, and *H* denote the population densities of the resident species, the invading species, and the hybrids, respectively. To evaluate the role of mating systems in shaping ecological outcomes, we first analyze a reproductive - interference scenario in which no hybrids are produced. We then extend the model to a backcrossing scenario in which viable hybrids can mate with either parent species.

The population dynamics for the parent species *i* (*i*, *j* = 1,2 and *i* ≠ *j*) are described as:

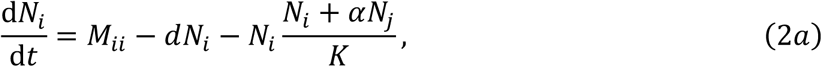

where *M_ii_* represent successful mating within species *i* as:

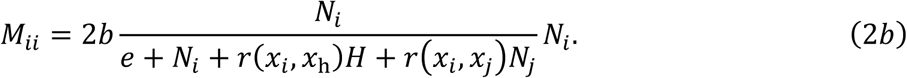

Model (ii): Backcrossing model

The backcrossing scenario is corresponding to the case when always *H* = 0 in Model (i). The population dynamics for the parent species *i* (*i*, *j* = 1,2 and *i* ≠ *j*) are described as:

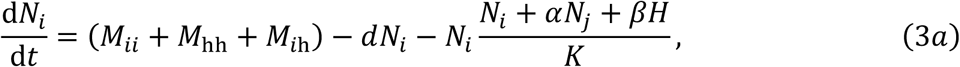

where the first term represents the sum of offspring from three types of successful matings. Specifically, *M_ii_* is the same term as Eq. (2b), and *M*_hh_ and *M*_*i*__h_ represent successful mating between hybrids, and between species *i* and hybrids, respectively:

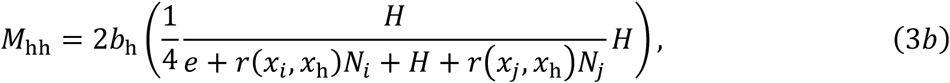

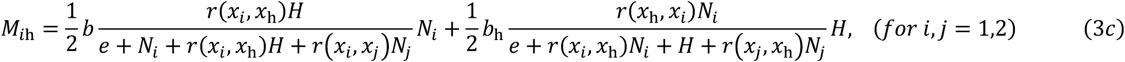

where *e* represents the cost of mate search. The number of successful matings for a female of type *i* with a male of type *j* is given by the birth rate of the female *b* multiplied by the proportion of males of type *j* she chooses (for *i*, *j* = 1,2, h and *i* ≠ *j*). This proportion (e.g., *M_ii_*) is determined by the mating probabilities and the abundance of all available males. It is noteworthy that offspring from *M*_hh_ and *M*_*i*__h_ matings can also result in *N_i_* individuals through Mendelian segregation. The former and latter terms of *M*_*i*__h_ represent matings between females of species *i* and hybrid males, and between males of species *i* and hybrid females, respectively. The second term describes mortality, with 𝑑*_i_* being the intrinsic death rate. The third term represents density-dependent mortality due to resource competition, where *K* is the carrying capacity, 𝛼 is the strength of interspecific competition between the two parent species, and 𝛽 is the competition strength between a parent species and the hybrids. Intraspecific competition is normalized to one. For simplicity, we assume symmetric competition and constant parameters.

We describe the population dynamics of hybrids as follows:

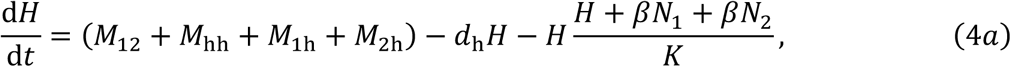

where 𝑑_h_ is the hybrid death rate. The hybrid offspring production term includes offspring from matings between the two parent species *M*_12_, between hybrids *M*_hh_, and between a parent species and a hybrid (*M*_1h_ and *M*_2h_). The number of offspring of a specific genotype resulting from each mating type is calculated based on Mendelian genetics, weighted by the female’s birth rate (*b* for parent species, 𝑏_h_ for hybrids). For example, the production of hybrid offspring (Aa) from matings between species 1 (AA) and species 2 (aa) is:

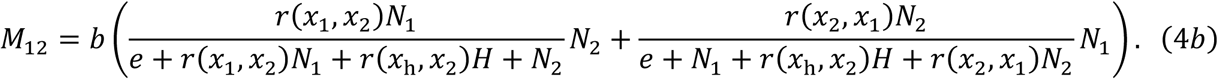

We can describe the matings within hybrids, *M*_hh_, those between species 1 and hybrid, *M*_1h_, and those between species 2 and hybrid, *M*_2h_.

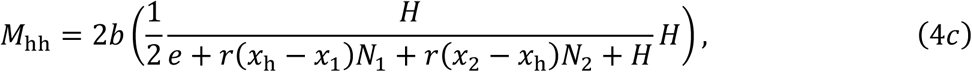

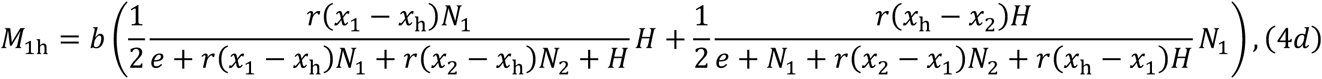

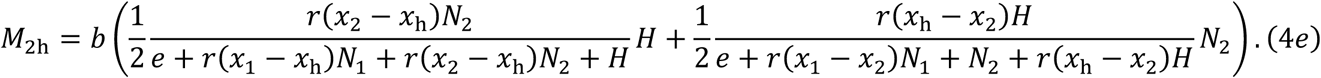

The fraction 1/2 in Eq. (3c) reflects the probability that hybrid genotypes are reproduced from hybrid × hybrid mating (Table S1). All computations for both models were performed using Mathematica (Wolfram 2021).

## 3. Result

We investigated how species coexistence is affected by the interplay between assortative mating, hybridization, and competition. We first analyzed a simplified scenario where hybrids are non-viable to isolate the effects of reproductive interference by analyzing Model (i). We then explored the full Model (ii) where viable hybrids can backcross with parent species, fundamentally altering the whole dynamics of the system.

### 3.1. Coexistence under reproductive interference without backcrossing

First, we considered the scenario of Model (i) where hybrids are assumed inviable. This simplification allows us to isolate the effects of reproductive interference. Using bifurcation diagrams, we assessed the stability of equilibria across a gradient of female choosiness, 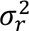.

When mate search is cost-free (i.e., *e* = 0), strong choosiness (i.e., low 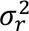) prevents reproductive interference and thus facilitates species coexistence. However, this coexistence is frequency-dependent; a species starting at a low density is always excluded. Weak mate preference (i.e., high 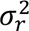) leads to the competitive exclusion of the rarer species, making coexistence impossible. These patterns are consistent across various levels of interspecific resource competition (Fig. 1a; Figs. S1c, e).

**Figure 1:**
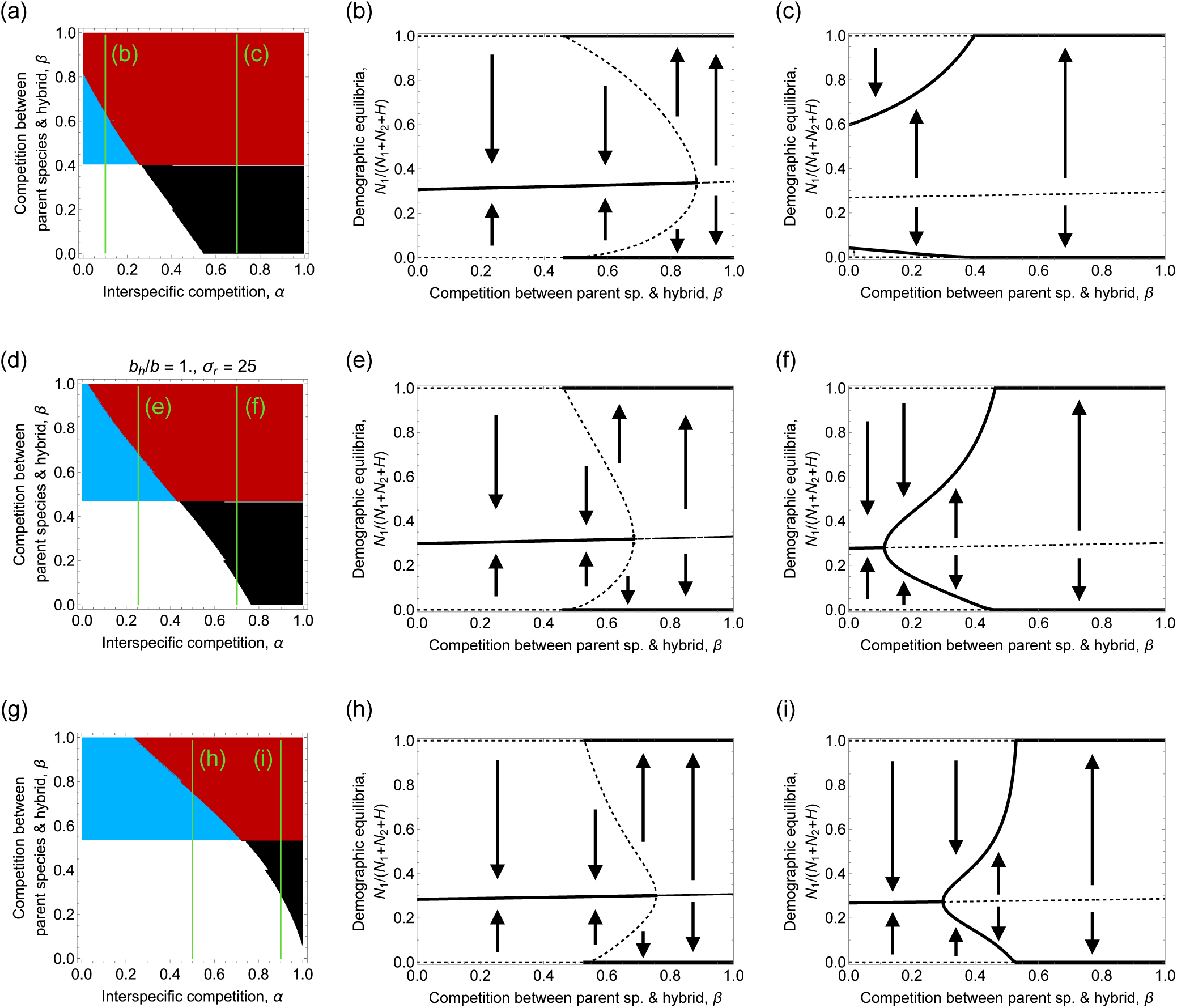
(a, d, g) The categories of ecological outcomes depending on the strengths of interspecific resource competition, 𝛼 and resource competition between parent species and hybrid, 𝛽 and (b, c, e, f, h, i) bifurcation diagrams of demographic equilibria of resident species 1. (a, d, g) We obtain the following four outcomes when searching for mating partner incurs no cost (i.e., *e* = 0): (i) alternative stable states (red), (ii) all-coexistence (white), (iii) frequency-dependent coexistence (the relative frequency of a rarer species determines coexistence) (blue), and (iv) competitive exclusion of rare species (red). Frequency-dependent coexistence indicates that if the initial densities of two species are similar, coexistence is possible, while a rare species goes extinct if one of the species has a sufficiently low density relative to its competitor species. (b, c, e, f, h, i) We show the frequency of resident species at equilibrium along the change in the strength of competition between parent species and hybrid. Solid and dashed lines denote locally stable and unstable equilibria, respectively. Parameters are 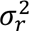 = 25, and 𝑏_h_ = (a) 0.75, (b) 1, (c) 1.25. Other parameters are 𝑏 = 1, 𝑑 = 𝑑_h_ = 0.1, 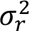 = 25, 𝑥_1_ = 0, 𝑥_2_ = 2, 𝑥_h_ = 1, and *K* = 1000.

Next, we focus on the scenario in which females incur a high cost of mate search (i.e., *e* = 450) to examine how this cost alters ecological outcomes compared to the case without such a cost. When a substantial cost of mate search is introduced, mate finding difficulty can drive both species to extinction, particularly when their initial densities are low (Figs. S1f–j). The reason is that an unstable equilibrium emerges such that, when the initial densities of both species fall below this point, the populations deterministically proceed toward extinction; this equilibrium thus represents an extinction threshold (the white circle near (50, 50) in Fig. S1d, e, i, and j). Thus, reproductive interference alone is sufficient to cause the competitive exclusion of the rare species, with outcomes depending on female mate choice and initial species densities.

### 3.2. Coexistence with viable hybrids and backcrossing

#### 3.2.1. Analytical conditions for species exclusion

We next considered the case of Model (ii) where hybrid offspring are viable and fertile, making the hybrid population density, *H*, a dynamic variable in Eqs. 2 and 3. This introduces the potential for backcrossing and competition with hybrids. For comparison with Scenario 1 (reproductive interference), we focused on the conditions of weak choosiness (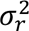 = 25), under which coexistence was impossible without viable hybrids.

We first analyzed the condition under which a single-species equilibrium (i.e., 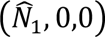 and 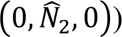 is locally stable––a rarer species is excluded competitively. Given that searching for mating partners has no cost (i.e., *e* = 0), the equilibria are locally stable when:

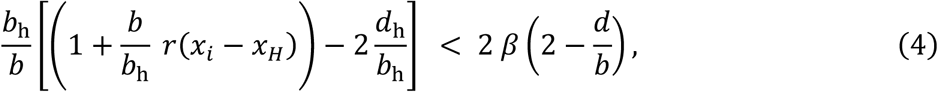

where 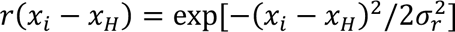. This inequality indicates that the emergence of the equilibria is independent of interspecific competition between parent species, 𝛼, but is dependent on female choosiness, 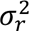, the birth and death rates of hybrids relative to those of the parent species, 𝑏/𝑏_h_ and 𝑑/𝑑_h_, and competition between parent species and hybrids, 𝛽 (see blue area in Figs. S2). Stronger parent-hybrid competition leads to the competitive exclusion (Figs. S2 and S3a–d). An increase in the birth rate of hybrids expands the parameter space of single-species equilibrium (Figs. S2a–d). Similarly, a decrease in the death rate of hybrids contracts it (Figs. S2e–g). Notably, our phase diagram shows a modest impact of female choosiness on the likelihood of competitive exclusion (Fig. S2 and S3), because hybrids are always reproduced through hybridization, backcrossing, and hybrid × hybrid mating regardless of the magnitude of female choosiness. Our system has minor effect of assortative mating on the outcome, which indicates that hybrid population dynamics dominates the whole dynamics of both species and their hybrid. Thus, ecological outcomes are primarily determined by parameters influencing the per capita growth rate of hybrids.

When mate search is costly, all-extinction equilibrium where both parent species and a hybrid population go extinct emerges if the mortality of all three populations (species 1, species 2, and hybrids) is positive (i.e., 𝑑 > 0 and 𝑑_h_ > 0), contrasting with cost-free scenarios where this equilibrium is always unstable.

#### 3.2.2. Confirmation of analytical conditions by simulations

Numerical simulations of Eqs. 2 and 3 confirmed the analytical predictions and revealed additional coexistence possibilities (Fig. S3). Without interspecific competition (i.e., 𝛼 = 0), single-species equilibria emerge under strong parent-hybrid competition, splitting into two categories: inevitable exclusion of rare species (red, Figs. S3e–h) and frequency-dependent coexistence where similar initial densities enable coexistence (blue, Figs. S3e–h). Increasing the hybrid birth rate expands the parameter space for frequency-dependent coexistence. When competition between parent species and hybrid is weak, the single-species equilibrium is unstable, and coexistence of two parent species and a hybrid population always occurs (we refer to ‘all-coexistence’ hereafter). When species interact through resource competition (i.e., 𝛼 = 0.6), outcomes differ: strong competition eliminates frequency-dependent coexistence, while weak competition generates alternative stable states—either species can dominate depending on initial density conditions (black, Figs. S3i–l).

#### 3.2.3. Ecological outcomes across competition gradients

We categorize ecological outcomes based on the strength of competition between species, 𝛼 and between parent species and hybrid, 𝛽 (Figs. 1 and 2). We search for the three-species-equilibria where all the densities of two species and a hybrid population are positive (i.e., 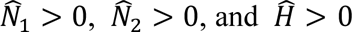) and the single-species equilibria (i.e., 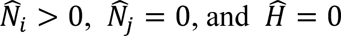), and numerically solve the local stability of the equilibria. Without the search cost (i.e., *e* = 0), we found four distinct outcomes independent of the relative rates of birth and death of hybrid over parent species (Fig. 1): (i) alternative stable states where two three-species-equilibria are locally stable and both single-species equilibria are unstable (black region in Fig. 1 and Fig. 2a), (ii) all-coexistence where one three-species-equilibrium is locally stable and both single-species equilibria are unstable (white region in Fig.1 and Fig. 2b), (iii) frequency-dependent coexistence where one three-species-equilibrium and both single-species equilibria are locally stable (blue region in Fig.1 and Fig. 2c), and (iv) competitive exclusion of a rarer species where all the three-species-equilibria are unstable and both single-species equilibria are locally stable (red region in Fig. 1 and Fig. 2d).

**Figure 2:**
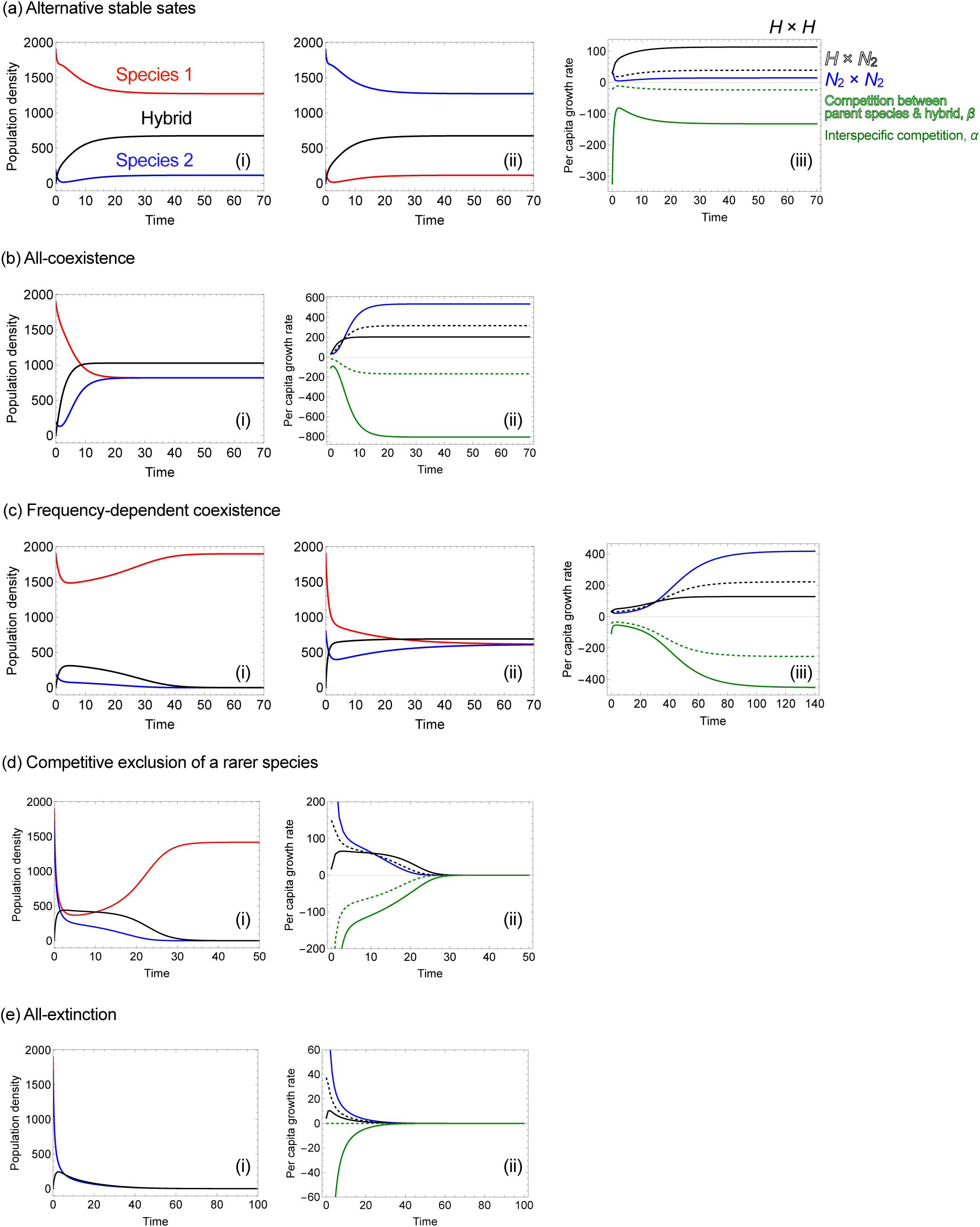
Population dynamics and changes in per capita growth rates under ecological outcomes. We first obtain the following four outcomes when searching for mating partner incurs no cost (*e* = 0). (a) Alternative stable states: (i) coexistence where species 1 (red) is dominant is possible, while (ii) coexistence is also possible when the initial density of species 2 (blue) is larger than that of species 1. (b) All-coexistence: coexistence of both species and their hybrid population is always possible. The equilibria of both species are equal. (c) Frequency-dependent coexistence: the initial frequency of species 2 determines coexistence. (i) a rare species goes extinct if one of the species has a sufficiently low density relative to its competitor species, while (ii) coexistence is possible if the initial densities of two species are similar. (d) Competitive exclusion of a rarer species always occurs. (e) All-extinction: when searching cost is high (*e* = 450), both species go extinct even when both species are initially abundant. We show the per capita growth rate of species 2 over time in panels (a–iii), (b–ii), (c–iii), (d–ii), and (e–ii). A blue solid line denotes the growth rate arising from conspecific mating and black solid and dashed lines denote the mating of hybrids and hybridization between invasive species and hybrid, respectively. Green solid and dashed denote the reduction of the growth rate due to competitive interactions, respectively. Parameters are (𝛼, 𝛽) = (a) (0.8,0.3), (b) (0.2,0.2), (c) (0.2,0.6), (d) (0.8,0.7), and (e) (0,0). Additionally, (a–d) 𝑏 = 𝑏_h_ = 1 and (e) 𝑏 = 𝑏_h_ = 0.25. The other parameters are 𝑑 = 𝑑_h_ = 0.1, 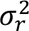 = 25, 𝑥 = 0, 𝑥 = 2, 𝑥 = (𝑥 + 𝑥)/2, and *K* = 1000. The initial densities are (*N*_1_, *N*_2_, *H*) = (a–c) ((2𝑏 − 𝑑)/*K*, 0.1 (2𝑏 − 𝑑)/*K*, 0’ and (d, e) ((2𝑏 − 𝑑)/*K*, 0.9 (2𝑏 − 𝑑)/*K*, 0).

Our numerical analysis confirms that the strength of resource competition between parent species and hybrid, 𝛽 determines the parameter space where single-species equilibria emerge (i.e., white/black regions versus blue/red regions in Fig. 1). In addition, the strength of interspecific resource competition, 𝛼 influences the emergence of equilibria where coexistence is possible. There are two cases of coexistence. One is that both parent species have the same densities (i.e., all-coexistence). As the other case, two equilibria––one where a resident species is more abundant than an invasive species and the other where the invasive one is more abundant––are feasible (i.e., alternative stable states; we show the underlying mechanism below). Strong interspecific competition 𝛼 shifts the system from all-coexistence (white region) to alternative stable states (black region) under weak 𝛽.

When the hybrid birth rate is lower or higher than that of the parent species (as compared to the case where hybrid and parent species share the same rate of birth), the qualitative outcomes remain similar (Fig. 2a, g). In most cases of all-coexistence or frequency-dependent coexistence, the hybrid population density exceeds that of the two species, forming a hybrid swarm (Figs. 2a, d, g; S4a, c–f). This occurs because hybrids are produced through more mating types—hybridization between the two species, hybrid × hybrid mating, and backcrossing with both species—than the parent species, which reproduce through only three mating types: conspecific mating, mating with hybrids, and hybrid × hybrid mating.

As detailed below, two key processes underlie the observed coexistence patterns: (i) regeneration of parental genotypes via hybridization, and (ii) competitive exclusion driven by hybrids. To clarify the mechanistic basis of coexistence patterns, we tracked temporal changes in mating contributions to per capita growth rate, *M_ij_* (*i*, *j* = 1,2, h) of Eqs. (2b), (3b), (3c) and (4b)–(4e) (within species 2, within hybrids, and between hybrids and species 2), as well as the effects of interspecific and hybrid–parent competition on per capita growth rate, −(*N_i_* + 𝛼*N_j_* + 𝛽*H*)/*K* (*i*, *j* = 1,2, *i* ≠ *j*) in Eq. (3a) and −(*H* + 𝛽*N*_1_ + 𝛽*N*_2_)/*K* in Eq. (4a) (Figs. 2a-iii, 2b-ii, 2c-iii, 2d-ii, 2e-ii). A key case of process (i) is shown in Fig. 2b-i: coexistence is achieved because hybrid matings regenerate parental genotypes through segregation. Despite reproductive interference preventing coexistence without hybrids, viable hybrids enable rare species recovery through genotype supply from hybrid populations. In contrast, strong competition between hybrids and parent species drives exclusion of the rare species (process (ii); Fig. 2d).

The balance between the two processes determines outcomes. Frequency-dependent coexistence occurs when they balance (Fig. 2c). Alternative stable states arise under strong interspecific but weak parent-hybrid competition (Fig. 2a). Strong interspecific competition excludes a rarer species, while weak parent-hybrid competition allows rare species to persist primarily through hybrid × hybrid mating. This is analogous to source–sink dynamics in metapopulation theory (Pulliam 1988). In Fig. 2a-iii, mating between hybrids (black line) exceeds that within species 2 (blue line).

With high mate search cost (i.e., *e* = 450), we basically obtained the same four outcomes (Fig. S5e-h) when the simulation starts with the resident species at carrying capacity. However, an additional outcome appears: both parent species and hybrids go extinct (Fig. 2e and Fig. S7e) through mate finding difficulty when the fitness of both species and hybrid is low, as our analysis showed that all-extinction equilibrium emerges in Result 3.2.1.

We evaluated the robustness of these coexistence categories by varying the birth and death rates of hybrids relative to those of the parent species. As the relative birth rate of hybrids increased, the coexistence-related parameter space expanded—regardless of the mate search cost (white and black areas in Fig. S5). Within the space, the space of all-coexistence became larger (white areas), but that of alternative stable states became narrower (black areas). On the other hand, as the rate increased, the extinction-related space contracted (blue and red areas in Fig. S5). Moreover, the space of frequency-dependent coexistence increased, while the space of competitive exclusion contracted. Similar patterns emerged with reduced hybrid mortality (Fig. S6).

We set the same birth rates among the three populations (𝑏 = 𝑏*ₕ*) and varied their values. The result revealed that high birth rates did not qualitatively alter the ecological outcomes, irrespective of search cost (Figs. S7b–d, f–h). In contrast, low birth rates increased the risk of extinction, particularly when mate search incurred a cost (Figs. S7a, e). Under high search costs, mate finding difficulty drove all-extinction (Fig. S7e).

## 4. Discussion

Our results underscore the importance of competitive interactions among parent species and their hybrids in shaping coexistence and exclusion outcomes in hybridizing systems. By explicitly coupling population dynamics with hybridization genetics, we identified five distinct ecological outcomes that emerge from the interplay between competition and backcrossing: (i) alternative stable states of coexistence, (ii) all-coexistence where both species and their hybrid population always persist, independent of their initial densities, (iii) frequency-dependent coexistence where coexistence is possible only when initial densities of two species are similar, (iv) competitive exclusion of a rarer species, and (v) all-extinction where both species and their hybrid go extinct. These outcomes arise from the interplay between two key processes: regeneration of parental genotypes through hybrid × hybrid mating, and competitive exclusion mediated by hybrids. Critically, our results show that backcrossing can enable coexistence even under competitive conditions that would otherwise lead to exclusion, revealing a previously underappreciated mechanism for maintaining species diversity in natural communities.

Historically, hybridization research has focused on its evolutionary consequences—especially its role in reinforcement and reproductive character displacement—because hybridization can either promote or hinder speciation (Liou and Price 1994, Goldberg and Lande 2006, Konuma and Chiba 2007, Mallet 2007, Abbott et al. 2013, Kyogoku and Yamaguchi 2023). Recent studies have highlighted its ecological consequences, including increased extinction risk when non-native species hybridize with resident species (Todesco et al. 2016, Porretta and Canestrelli 2023). Even under simplified assumptions about hybridization genetics, our model shows that multiple coexistence regimes emerge from eco-genetic interactions involving competitive dynamics among parent species and hybrids, rather than from hybridization genetics alone.

Previous empirical studies have mainly focused on factors related to hybrid fitness and reproductive isolation (e.g., Rieseberg et al. (2003) and Taylor and Larson (2019)). In our model, hybrid fitness is represented by the rates of birth and death, and we demonstrated a novel result: the four outcomes of coexistence and extinction (all-coexistence, frequency-dependent coexistence, alternative stable states of coexistence, and competitive exclusion of a rare species) are robust and occur independently of hybrid fitness. While theoretical work traditionally assumes ecological neutrality between hybridizing species, empirical studies show that competition coefficients between parent species (𝛼) and between parents and hybrids (𝛽) vary widely across taxa. For instance, closely related species often experience strong competition depending on niche and competitive-ability differences (Chesson 2000, Levine and HilleRisLambers 2009), and hybrids show stronger competitive ability than their parent species in several taxa, leading to larger 𝛽 values (Boersma 1995, Seiler and Keeley 2007). Despite their importance, empirical knowledge about the relative strengths of 𝛼 and 𝛽 remains very limited (e.g., Freeland et al. (2025)). Our results reveal that these relative strengths of 𝛼 and 𝛽 significantly affect the coexistence of parent species. Specifically, our model predicts that when competition between parent species and hybrid is stronger than that between parent species (𝛼 < 𝛽), coexistence becomes frequency dependent, making initial relative densities a decisive factor.

The balance between 𝛼 and 𝛽 is closely tied to the niche traits expressed in hybrids, thereby shaping their potential to colonize either intermediate or novel ecological space. We interpret 𝛼 and 𝛽 such that hybrid niche traits modify the relative strength of parent–hybrid competition compared with interspecific competition. When 𝛼 < 𝛽, hybrids typically occupy an intermediate niche between their parent species (Abbott et al. 2013; see example of two niche traits in Fig. S8a). Such patterns have been empirically documented in a wide range of plant and animal taxa, such as *Heliconius* butterflies (Mallet et al. 2007). Although intermediate niches can maintain stable hybrid zones, they rarely foster speciation because continuous contact and backcrossing with parent species erode lineage independence (Barton and Hewitt 1989, Harrison and Larson 2014). By contrast, when 𝛼 > 𝛽, hybrids can establish in a novel niche outside that of parent species (see example in Fig. S8b), a scenario with profound evolutionary significance. Such ecological novelty often arises through transgressive segregation, in which recombination generates trait values beyond the parental range via complementary additive alleles (Rieseberg 1997, Stelkens and Seehausen 2009, Yamaguchi and Otto 2020). This process represents a potential mechanism for homoploid hybrid speciation, speciation without a change in ploidy (Rieseberg 1997), as shown in several *Helianthus* species in North America (Rieseberg et al. 1990), where hybrid-origin lineages successfully colonized extreme environments distinct from those of their parental species. However, homoploid hybrid speciation remains rare in nature, requiring a unique combination of reproductive isolation and ecological opportunity (Mallet 2007, Schumer et al. 2014).

Two of the five regimes predicted by our model—stable all-coexistence and competitive exclusion of the rarer species—reflects patterns frequently observed in natural hybrid zones. For instance, some hybrid zones, such as those in *Senecio* plants, are known to be stably maintained for long periods (e.g., Abbott et al. (2003), Brennan et al. (2013)), corresponding to our “all-coexistence” scenario. Conversely, the rapid displacement of a native species by invasive hybrids, as seen in sunfish and trout, provides clear evidence for competitive exclusion (e.g., Muhlfeld et al. (2009)). The outcome of frequency-dependent coexistence suggests that the history of colonization can be critical; one could potentially find two distinct geographical locations involving the same two species where one forms a stable hybrid zone while the other experiences displacement, reflecting the different colonizing history. More importantly, our model offers a novel, intrinsic mechanism for the formation of “mosaic” hybrid zones through the “alternative stable states of coexistence.” Traditionally, mosaic hybrid zones have been attributed to species sorting across environmentally heterogeneous landscapes (Harrison 1986, Ross and Harrison 2002), while our model predicts that such a biogeographical pattern can emerge even in a homogeneous environment. In this scenario, the dominance of either parent species in a given patch is determined solely by initial densities and the dynamics of genotype regeneration, with immigration from adjacent patches.

Predictions involving all-extinction are difficult to verify in the field, yet they highlight scenarios where hybridization depresses densities so strongly that mate finding difficulty triggers local extinction—a process likely underappreciated in hybridizing systems.

Our model also helps explain empirical contrasts between secondary contact and biological invasion. Secondary contact typically involves species meeting at similar densities, whereas invasions begin with rare immigrants entering established populations. Because our model reveals strong frequency dependence, these different initial demographic contexts generate distinct outcomes: stable coexistence under secondary contact versus genetic swamping or exclusion under invasion (Levin et al. 1993, Rhymer and Simberloff 1996, Todesco et al. 2016). For example, in sunflowers (*Helianthus* spp.), secondary contact between closely related species has produced long-term hybrid zones where both parent species persist (Rieseberg et al. 2007). By contrast, in waterfowl such as mallards (*Anas platyrhynchos*) invading the ranges of native ducks, extensive hybridization has led to competitive exclusion of rare native species (Rhymer and Simberloff 1996). Our results provide a mechanistic explanation for these contrasting outcomes through frequency dependence in ecological communities, consistent with priority effects (Fukami 2015). Thus, our model highlights the importance of the history of contact between hybridizing species as a key determinant of hybrid zone dynamics.

Although challenging to detect in the field, this prediction is highly testable under controlled experimental conditions. Manipulative mesocosm or common garden studies that vary initial densities and quantify competition parameters could provide direct evidence of whether hybridization drives coexistence, exclusion, or collapse. Integrating such experimental tests with field observations of natural hybrid zones would greatly advance our ability to evaluate the ecological consequences of hybridization.

Previous theoretical studies have primarily attributed species extinction in hybridizing systems to genetic mechanisms alone (Huxel 1999, Wolf et al. 2001, Ferdy and Austerlitz 2002, Thompson et al. 2003, Hall and Ayres 2009). Our study broadens this paradigm by identifying two interacting mechanisms—genotype regeneration and hybrid-mediated competitive exclusion—as central determinants of coexistence regimes. Consistent with earlier work, our model reproduces genetic displacement outcomes, particularly under high hybrid birth and death rates (i.e., heterosis) (Huxel 1999, Wolf et al. 2001).

Among the few studies incorporating both genetic and ecological processes, our model not only encompasses but also complements existing approaches. For example, Hall et al. (2006) showed that hybrid’s reproductive traits, such as fecundity, strongly shape parent–hybrid dynamics, but did not explicitly model competitive interactions between parent species and their hybrid. Our results partially align with those of Irwin and Schluter (2022), yet we uncover additional coexistence regimes and the modest effects of assortative mating. These differences arise from fundamental differences in genetic architecture: in polygenic models, such as that of Irwin and Schluter (2022), genotype regeneration is relatively unlikely, leading to blending into a hybrid population, whereas our Mendelian framework allows parental genotypes to re-emerge. This distinction suggests that genetic architecture—specifically the potential for genotype regeneration—may be crucial for understanding species persistence in hybrid zones. Although previous polygenic models have shown that genotype regeneration can contribute to genetic or evolutionary rescue (Mesgaran et al. 2016, Vedder et al. 2022), it remains necessary to test whether qualitatively similar results emerge when implementing an individual-based polygenic model. By integrating competitive dynamics into a genetically explicit framework, our Lotka-Volterra-based model provides a tractable platform for unifying population genetics and ecology in hybridization studies, addressing the broader need for eco-evolutionary synthesis (Lambrinos 2004).

Our model relies on several simplifying assumptions that open avenues for future research. First, we modeled reproductive interference through incomplete mate choice, whereas other mechanisms—such as heterospecific mating harassment—may lead to different outcomes (Gröning and Hochkirch 2008, Kawatsu 2013). Also, speciation models sometimes consider two types of traits explicitly––female choosiness and ecological selected traits––in addition to mating signals and female preference our model (Doebeli 2005, Kopp et al. 2018). Incorporating alternative interference processes and recombination processes of additional traits could reveal additional coexistence mechanisms. Second, we assumed symmetric and constant competition coefficients, as well as constant birth and death rates, despite evidence that asymmetric interspecific competition is common (Weiner 1990, Vázquez et al. 2007, Violle et al. 2012). Examining asymmetric competition or incorporating density-dependent demographic rates could alter the predicted stability of local coexistence. Finally, our model lacks spatial structure, despite the documented role of spatial heterogeneity in shaping hybrid zones (Buggs 2007, Hall and Ayres 2009, Brennan et al. 2015) and theoretical predictions that spatial expansion dynamics can fundamentally alter hybrid fitness landscapes (MacPherson et al. 2022). Incorporating spatial dynamics would generate localized coexistence patterns and spatial refugia that prevent global extinction.

In conclusion, our study shows that hybridization does not inevitably threaten species persistence. Instead, it can either stabilize coexistence or precipitate exclusion, depending on the competitive relationships among parent species and their hybrids. Whereas previous work has emphasized genetic mechanisms driving extinction under hybridization, our analysis reveals that genotype regeneration through hybrid × hybrid mating can promote long-term coexistence. Thus, hybridization operates as a double-edged sword, capable of both maintaining and collapsing ecological communities. By highlighting how ecological competition and hybridization genetics interact to generate multiple coexistence regimes, our results underscore the need for future research to move beyond genetically focused frameworks and incorporate ecological dynamics more explicitly. Such integration will be essential for predicting the diverse ecological outcomes of hybridizing species in a rapidly changing world.

## Authors’ contributions

KM: Data curation, Data curation, Formal analysis, Methodology, Project administration, Software, Visualization, and Writing – original draft

RY: Funding acquisition, Methodology, and Writing – review & editing

## Declaration of AI use

Some sections of the manuscript were edited for grammar and style using ChatGPT (OpenAI, 2025). The authors reviewed, corrected, and approved all content, and take full responsibility for its accuracy.

## Acknowledgement

Dr. Daisuke Kyogoku gave us helpful comments.

## Reference

Abbott, R. J., James, J. K., Milne, R. I. and Gillies, A. C. M. 2003. Plant introductions, hybridization and gene flow (HG Dickinson, SJ Hiscock, and PR Crane, Eds.). - Phil. Trans. R. Soc. Lond. B 358: 1123‒1132.

Abbott, R., Albach, D., Ansell, S., Arntzen, J. W., Baird, S. J. E., Bierne, N., Boughman, J., Brelsford, A., Buerkle, C. A., Buggs, R., Butlin, R. K., Dieckmann, U., Eroukhmanoff, F., Grill, A., Cahan, S. H., Hermansen, J. S., Hewitt, G., Hudson, A. G., Jiggins, C., Jones, J., Keller, B., Marczewski, T., Mallet, J., Martinez-Rodriguez, P., Möst, M., Mullen, S., Nichols, R., Nolte, A. W., Parisod, C., Pfennig, K., Rice, A. M., Ritchie, M. G., Seifert, B., Smadja, C. M., Stelkens, R., Szymura, J. M., Väinölä, R., Wolf, J. B. W. and Zinner, D. 2013. Hybridization and speciation. - J of Evolutionary Biology 26: 229‒246.

Ålund, M., Marzal, J. C. S., Zhu, Y., Menon, P. N. K., Jones, W. and Qvarnström, A. 2024. Tracking hybrid viability across life stages in a natural avian contact zone (S Taylor and T Connallon, Eds.). - Evolution 78: 267‒283.

Arntzen, J. W., De Vries, W., Canestrelli, D. and Martínez - Solano, I. 2017. Hybrid zone formation and contrasting outcomes of secondary contact over transects in common toads. - Molecular Ecology 26: 5663‒5675.

Ayres, D. R., Zaremba, K. and Strong, D. R. 2004. Extinction of a Common Native Species by Hybridization with an Invasive Congener^1^. - Weed Technology 18: 1288‒1291.

Barton, N. H. and Hewitt, G. M. 1989. Adaptation, speciation and hybrid zones. - Nature 341: 497‒503.

Boersma, M. 1995. Competition in natural populations of Daphnia. - Oecologia 103: 309‒318.

Brennan, A. C., Harris, S. A. and Hiscock, S. J. 2013. THE POPULATION GENETICS OF SPOROPHYTIC SELF-INCOMPATIBILITY IN THREE HYBRIDIZING *SENECIO* (ASTERACEAE) SPECIES WITH CONTRASTING POPULATION HISTORIES: SELF-INCOMPATIBILITY AND HYBRIDIZATION. - Evolution: no-no.

Brennan, A. C., Woodward, G., Seehausen, O., Muñoz-Fuentes, V., Moritz, C., Guelmami, A., Abbott, R. J. and Edelaar, P. 2015. Hybridization due to changing species distributions: adding problems or solutions to conservation of biodiversity during global change? - Evolutionary Ecology Research 16: 475‒491.

Buggs, R. J. A. 2007. Empirical study of hybrid zone movement. - Heredity 99: 301‒312.

Chesson, P. 2000. General Theory of Competitive Coexistence in Spatially-Varying Environments. - Theoretical Population Biology 58: 211‒237.

Doebeli, M. 2005. Adaptive speciation when assortative mating is based on female preference for male marker traits. - Journal of Evolutionary Biology 18: 1587‒1600.

Donovan, L. A., Rosenthal, D. R., Sanchez - Velenosi, M., Rieseberg, L. H. and Ludwig, F. 2010. Are hybrid species more fit than ancestral parent species in the current hybrid species habitats? - J of Evolutionary Biology 23: 805‒816.

Epifanio, J. and Philipp, D. 2000. Simulating the extinction of parental lineages from introgressive hybridization: the effects of fitness, initial proportions of parental taxa, and mate choice. - Reviews in Fish Biology and Fisheries 10: 339‒354.

Ferdy, J. and Austerlitz, F. 2002. Extinction and Introgression in a Community of Partially Cross - Fertile Plant Species. - The American Naturalist 160: 74‒86.

Freeland, J., Kowalcyk, O., Brennan, M. and Dorken, M. 2025. Quality not Quantity: Seedlings of the Invasive Hybrid Cattail Typha x glauca Outcompete the more Abundant Seedlings of their Maternal Parent T. angustifolia. - Wetlands 45: 6.

Fukami, T. 2015. Historical Contingency in Community Assembly: Integrating Niches, Species Pools, and Priority Effects. - Annu. Rev. Ecol. Evol. Syst. 46: 1‒23.

Goldberg, E. and Lande, R. 2006. Ecological and reproductive character displacement of an environmental gradient. - Evolution 60: 1344‒1357.

Gröning, J. and Hochkirch, A. 2008. Reproductive Interference Between Animal Species. - The Quarterly Review of Biology 83: 257‒282.

Hall, R. J. and Ayres, D. R. 2009. What can mathematical modeling tell us about hybrid invasions? - Biol Invasions 11: 1217‒1224.

Hall, R. J., Hastings, A. and Ayres, D. R. 2006. Explaining the explosion: modelling hybrid invasions. - Proc. R. Soc. B. 273: 1385‒1389.

Harrison, R. G. 1986. Pattern and process in a narrow hybrid zone. - Heredity 56: 337‒349.

Harrison, R. G. and Larson, E. L. 2014. Hybridization, introgression, and the nature of species boundaries. - Journal of Heredity 105: 795‒809.

Hauber, M. E. and Sherman, P. W. 2001. Self-referent phenotype matching: theoretical considerations and empirical evidence. - Trends in neurosciences 24: 609‒616.

Huxel, G. R. 1999. Rapid displacement of native species by invasive species: effects of hybridization. - Biological Conservation 89: 143‒152.

Irwin, D. and Schluter, D. 2022. Hybridization and the Coexistence of Species. - The American Naturalist 200: E93‒E109.

Kawatsu, K. 2013. Sexually Antagonistic Coevolution for Sexual Harassment Can Act as a Barrier to Further Invasions by Parthenogenesis. - The American Naturalist 181: 223‒234.

Kishi, S. and Nakazawa, T. 2013. Analysis of species coexistence co - mediated by resource competition and reproductive interference. - Population Ecology 55: 305‒313.

Konuma, J. and Chiba, S. 2007. Ecological character displacement caused by reproductive interference. - Journal of theoretical Biology 247: 354‒364.

Kopp, M., Servedio, M. R., Mendelson, T. C., Safran, R. J., Rodríguez, R. L., Hauber, M. E., Scordato, E. C., Symes, L. B., Balakrishnan, C. N., Zonana, D. M. and Van Doorn, G. S. 2018. Mechanisms of Assortative Mating in Speciation with Gene Flow: Connecting Theory and Empirical Research. - The American Naturalist 191: 1‒20.

Kuno, E. 1992. Competitive exclusion through reproductive interference. - Population Ecology 34: 275‒284.

Kyogoku, D. 2020. When does reproductive interference occur? Predictions and data. - Population Ecology 62: 196‒206.

Kyogoku, D. and Yamaguchi, R. 2023. Males and females contribute differently to the evolution of habitat segregation driven by hybridization. - j. evol. Biol. 36: 515‒528.

Lambrinos, J. G. 2004. HOW INTERACTIONS BETWEEN ECOLOGY AND EVOLUTION INFLUENCE CONTEMPORARY INVASION DYNAMICS. - Ecology 85: 2061‒2070.

Levin, D. A., Francisco - Ortega, J. and Jansen, R. K. 1996. Hybridization and the Extinction of Rare Plant Species. - Conservation Biology 10: 10‒16.

Levine, J. M. and HilleRisLambers, J. 2009. The importance of niches for the maintenance of species diversity. - Nature 461: 254‒257.

Liou, L. W. and Price, T. D. 1994. SPECIATION BY REINFORCEMENT OF PREMATING ISOLATION. - Evolution 48: 1451‒1459.

MacPherson, A., Wang, S., Yamaguchi, R., Rieseberg, L. H. and Otto, S. P. 2022. Parental Population Range Expansion before Secondary Contact Promotes Heterosis. - The American Naturalist 200: E1‒E15.

Mallet, J. 2007. Hybrid speciation. - Nature 446: 279‒283.

Mallet, J., Beltrán, M., Neukirchen, W. and Linares, M. 2007. Natural hybridization in heliconiine butterflies: the species boundary as a continuum. - BMC Evol Biol 7: 28.

Mandeville, E. G., Walters, A. W., Nordberg, B. J., Higgins, K. H., Burckhardt, J. C. and Wagner, C. E. 2019. Variable hybridization outcomes in trout are predicted by historical fish stocking and environmental context. - Molecular Ecology 28: 3738‒3755.

Mandeville, E. G., Hall, R. O. and Buerkle, C. A. 2022. Ecological outcomes of hybridization vary extensively in *Catostomus* fishes. - Evolution 76: 2697‒2711.

Martins, A. R. P., Warren, N. B., McMillan, W. O. and Barrett, R. D. H. 2024. Spatiotemporal dynamics in butterfly hybrid zones. - Insect Science 31: 328‒353.

Mesgaran, M. B., Lewis, M. A., Ades, P. K., Donohue, K., Ohadi, S., Li, C. and Cousens, R. D. 2016. Hybridization can facilitate species invasions, even without enhancing local adaptation. - Proc. Natl. Acad. Sci. U.S.A. 113: 10210‒10214.

Mittelbach, G. G. and McGill, B. J. 2019. Community ecology. - Oxford University Press.

Muhlfeld, C. C., Kalinowski, S. T., McMahon, T. E., Taper, M. L., Painter, S., Leary, R. F. and Allendorf, F. W. 2009. Hybridization rapidly reduces fitness of a native trout in the wild. - Biol. Lett. 5: 328‒331.

Pfennig, K. S., Kelly, A. L. and Pierce, A. A. 2016. Hybridization as a facilitator of species range expansion. - Proc. R. Soc. B. 283: 20161329.

Porretta, D. and Canestrelli, D. 2023. The ecological importance of hybridization. - Trends in Ecology & Evolution 38: 1097‒1108.

Prentis, P. J., White, E. M., Radford, I. J., Lowe, A. J. and Clarke, A. R. 2007. Can hybridization cause local extinction: a case for demographic swamping of the Australian native *Senecio pinnatifolius* by the invasive *Senecio madagascariensis* ? - New Phytologist 176: 902‒912.

Quilodrán, C. S., Austerlitz, F., Currat, M. and Montoya-Burgos, J. I. 2018. Cryptic Biological Invasions: a General Model of Hybridization. - Sci Rep 8: 2414.

Reed, T. E., Kane, A., McGinnity, P. and O’Sullivan, R. J. 2024. Competitive interactions affect introgression and population viability amidst maladaptive hybridization. - Evolutionary Applications 17: e13746.

Rhymer, J. M. and Simberloff, D. 1996. EXTINCTION BY HYBRIDIZATION AND INTROGRESSION. - Annu. Rev. Ecol. Syst. 27: 83‒109.

Rieseberg, L. H. 1997. Hybrid Origins of Plant Species. - Annu. Rev. Ecol. Syst. 28: 359‒389.

Rieseberg, L. H., Carter, R. and Zona, S. 1990. MOLECULAR TESTS OF THE HYPOTHESIZED HYBRID ORIGIN OF TWO DIPLOID *HELIANTHUS* SPECIES (ASTERACEAE). - Evolution 44: 1498‒1511.

Rieseberg, L. H., Raymond, O., Rosenthal, D. M., Lai, Z., Livingstone, K., Nakazato, T., Durphy, J. L., Schwarzbach, A. E., Donovan, L. A. and Lexer, C. 2003. Major Ecological Transitions in Wild Sunflowers Facilitated by Hybridization. - Science 301: 1211‒1216.

Rieseberg, L. H., Kim, S.-C., Randell, R. A., Whitney, K. D., Gross, B. L., Lexer, C. and Clay, K. 2007. Hybridization and the colonization of novel habitats by annual sunflowers. - Genetica 129: 149‒165.

Ross, C. L. and Harrison, R. G. 2002. A FINE-SCALE SPATIAL ANALYSIS OF THE MOSAIC HYBRID ZONE BETWEEN GRYLLUS FIRMUS AND GRYLLUS PENNSYLVANICUS. - Evolution 56: 2296‒2312.

Schierenbeck, K. A. and Ellstrand, N. C. 2009. Hybridization and the evolution of invasiveness in plants and other organisms. - Biol Invasions 11: 1093‒1105.

Schreiber, S. J., Yamamichi, M. and Strauss, S. Y. 2019. When rarity has costs: coexistence under positive frequency - dependence and environmental stochasticity. - Ecology 100: e02664.

Schumer, M., Rosenthal, G. G. and Andolfatto, P. 2014. HOW COMMON IS HOMOPLOID HYBRID SPECIATION?: PERSPECTIVE. - Evolution 68: 1553‒1560.

Seiler, S. M. and Keeley, E. R. 2007. A comparison of aggressive and foraging behaviour between juvenile cutthroat trout, rainbow trout and F1 hybrids. - Animal Behaviour 74: 1805‒1812.

Stelkens, R. and Seehausen, O. 2009. GENETIC DISTANCE BETWEEN SPECIES PREDICTS NOVEL TRAIT EXPRESSION IN THEIR HYBRIDS. - Evolution 63: 884‒897.

Taylor, S. A. and Larson, E. L. 2019. Insights from genomes into the evolutionary importance and prevalence of hybridization in nature. - Nat Ecol Evol 3: 170‒177.

Thompson, C. J., Thompson, B. J. P., Ades, P. K., Cousens, R., Garnier-Gere, P., Landman, K., Newbigin, E. and Burgman, M. A. 2003. Model-based analysis of the likelihood of gene introgression from genetically modified crops into wild relatives. - Ecological Modelling 162: 199‒209.

Todesco, M., Pascual, M. A., Owens, G. L., Ostevik, K. L., Moyers, B. T., Hübner, S., Heredia, S. M., Hahn, M. A., Caseys, C., Bock, D. G. and Rieseberg, L. H. 2016. Hybridization and extinction. - Evolutionary Applications 9: 892‒908.

Vallin, N., Rice, A. M., Arntsen, H., Kulma, K. and Qvarnström, A. 2012. Combined effects of interspecific competition and hybridization impede local coexistence of Ficedula flycatchers. - Evol Ecol 26: 927‒942.

Vázquez, D. P., Melián, C. J., Williams, N. M., Blüthgen, N., Krasnov, B. R. and Poulin, R. 2007. Species abundance and asymmetric interaction strength in ecological networks. - Oikos 116: 1120‒1127.

Vedder, D., Lens, L., Martin, C. A., Pellikka, P., Adhikari, H., Heiskanen, J., Engler, J. O. and Sarmento Cabral, J. 2022. Hybridization may aid evolutionary rescue of an endangered East African passerine. - Evolutionary Applications 15: 1177‒1188.

Violle, C., Enquist, B. J., McGill, B. J., Jiang, L., Albert, C. H., Hulshof, C., Jung, V. and Messier, J. 2012. The return of the variance: intraspecific variability in community ecology. - Trends in Ecology & Evolution 27: 244‒252.

Weiner, J. 1990. Asymmetric competition in plant populations. - Trends in Ecology & Evolution 5: 360‒364.

Williams, J., Lambert, A. M., Long, R. and Saltonstall, K. 2019. Does hybrid *Phragmites australis* differ from native and introduced lineages in reproductive, genetic, and morphological traits? - American J of Botany 106: 29‒41.

Wolf, D. E., Takebayashi, N. and Rieseberg, L. H. 2001. Predicting the Risk of Extinction through Hybridization. - Conservation Biology 15: 1039‒1053.

Yamaguchi, R. and Otto, S. P. 2020. Insights from Fisher’s geometric model on the likelihood of speciation under different histories of environmental change. - Evolution 74: 1603‒1619.

Yamaguchi, R., Yamanaka, T. and Liebhold, A. M. 2019. Consequences of hybridization during invasion on establishment success. - Theor Ecol 12: 197‒205.

