## Supporting Information for "Is hybridization evil? –– Competition generates alternative stable states of coexistence of hybridizing species"

**SUPPLEMENTARY FIGURE 1** 2

**SUPPLEMENTARY FIGURE 2** 3

**SUPPLEMENTARY FIGURE 3** 4

**SUPPLEMENTARY FIGURE 4** 5

**SUPPLEMENTARY FIGURE 5** 7

**SUPPLEMENTARY FIGURE 6** 8

**SUPPLEMENTARY FIGURE 7** 9

**SUPPLEMENTARY FIGURE 8** 10

**SUPPLEMENTARY TABLE** 11

### Supplementary figure 1

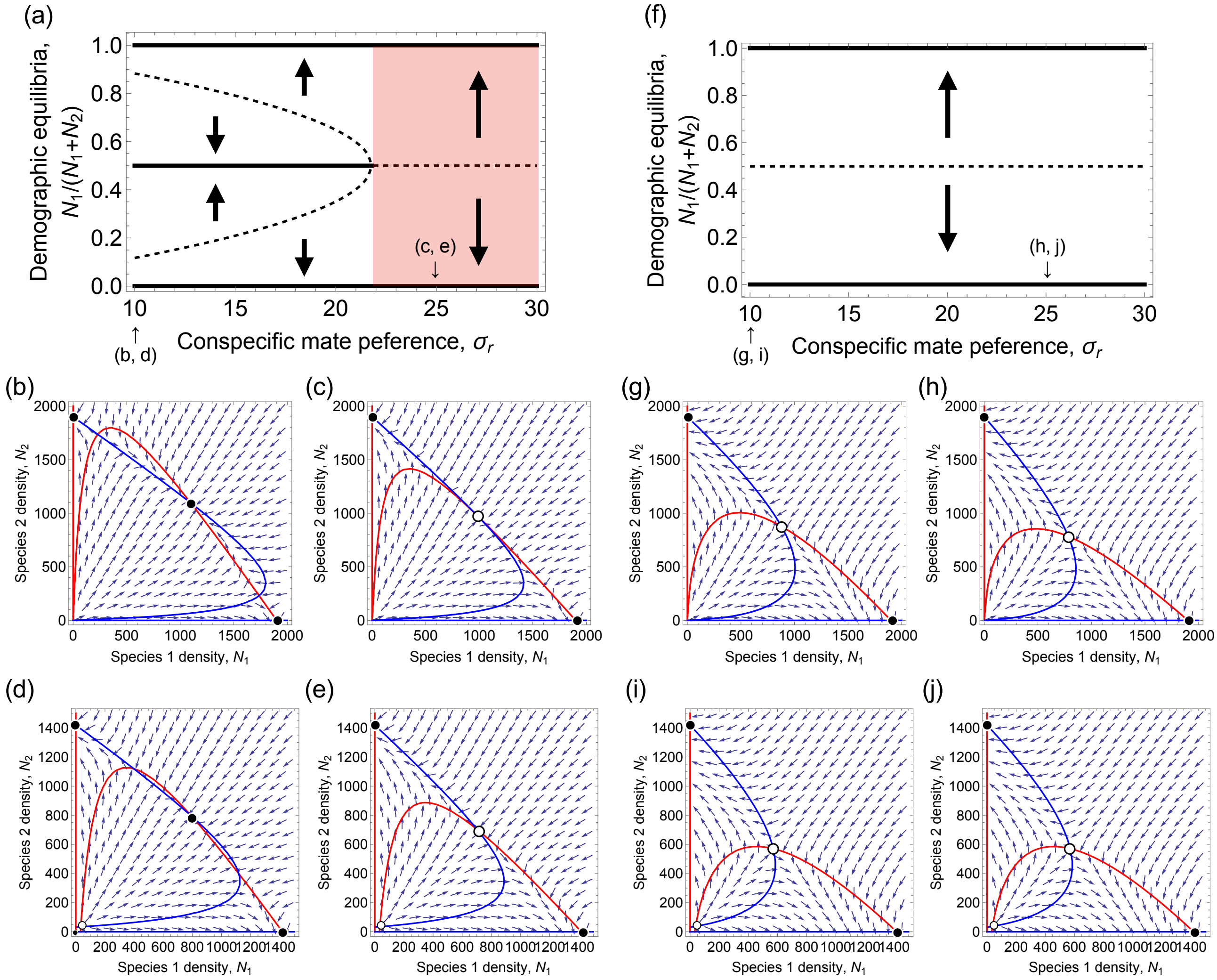


**Figure S1**: (a, f) Bifurcation diagrams and (b–e, g–j) nullcline plots of demographic equilibria of two species. When searching for mating partners incurs no cost ($e = 0$) and two species interact without resource competition ($\alpha = 0$), the rare interbreeding of two species leads to coexistence (b, $\sigma_r^2 = 10$), while the frequent interbreeding of two species drives rare species to extinction (c, $\sigma_r^2 = 25$). These results are qualitatively the same when two species interact with resource competition ($\alpha = 0.1$; d, $\sigma_r^2 = 10$; e, $\sigma_r^2 = 10$). When the searching is costly ($e = 450$), the extinction of rare species occurs regardless of the likelihood of the interbreeding (g, $\sigma_r^2 = 10$; h, $\sigma_r^2 = 25$) without resource competition. In the presence of resource competition, the extinction is more likely (i, $\sigma_r^2 = 10$; j, $\sigma_r^2 = 25$). Solid and dashed lines denote locally stable and unstable equilibria of the frequency of resident species 1, $N_1/(N_1 + N_2)$, respectively. (b–e, g–j) Nullcline plots of resident species (red) and invasive species (blue). Black and white points denote locally stable and unstable equilibria, respectively. Other parameters are $b = 1$, $d = 0.1$, $x_1 = 0$, $x_2 = 2$, and $K = 1000$.

### Supplementary figure 2

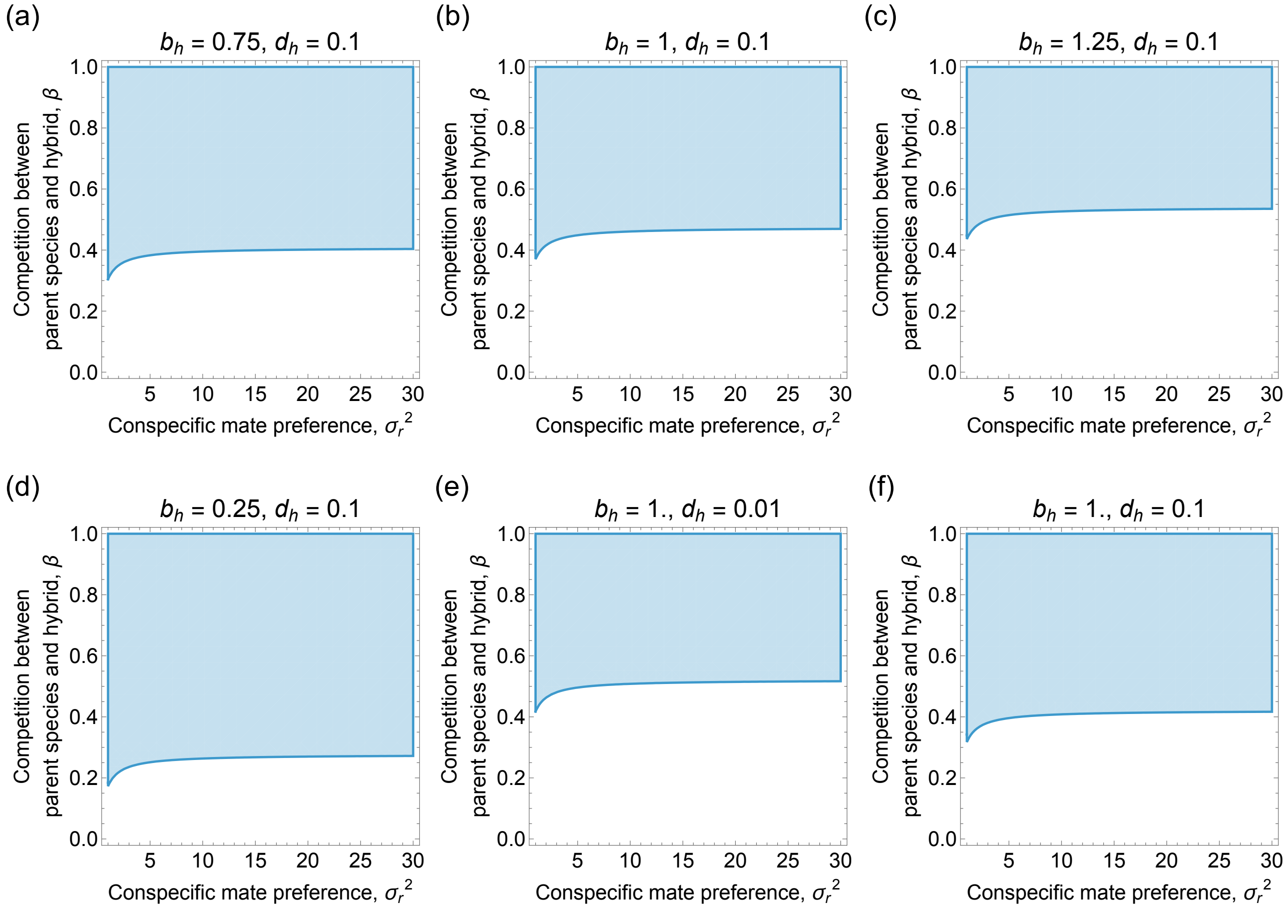


**Figure S2**: Analysis of local stability of the equilibria where the extinction of either species occurs when searching for mating partners incurs no cost ($e = 0$). Competitive exclusion (i.e., $(\hat{N}_1, \hat{N}_2, \hat{H}) = ((2b - d)/K, 0, 0), (0, (2b - d)/K, 0))$ occurs in blue areas. Parameters are $b_{\mathrm{h}} =$ (a) 0.75, (b) 1, (c) 1.25, and (d) 0.25 when the mortalities of parent species and hybrid are symmetries ($d_{\mathrm{h}} = d = 0.1$), while $d_{\mathrm{h}} =$ (e) 0.01 and (f) 0.2 when the birth rates of parent species and hybrid are symmetries (i.e., $b_{\mathrm{h}} = b = 1$). Other parameters are $\sigma_r^2 = 25$, $x_1 = 0$, $x_2 = 2$, $x_{\mathrm{h}} = 1$, and $K = 1000$.

### Supplementary figure 3

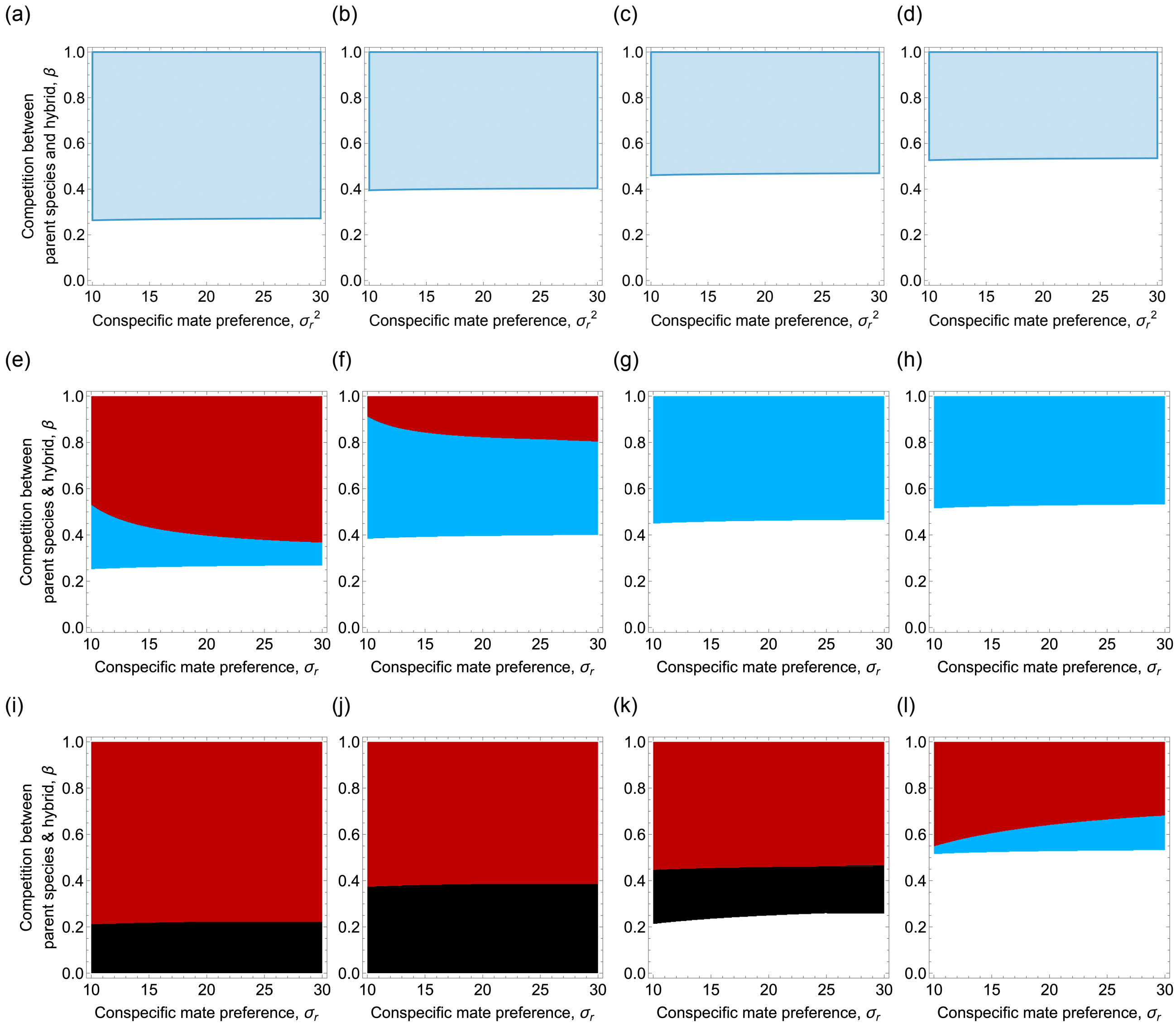


**Figure S3**: Competitive exclusion of either species is possible depending on the ratio of hybrid-to-parent-species birth rates and competition between parent species and hybrid when searching for mating partners incurs no cost ($e = 0$). (a-d) Analytical results when the hybrid birth rates are $b_{\mathrm{h}} =$ (a) 0.25, (b) 0.75, (c) 1, and (d) 1.25. Competitive exclusion (i.e., $(\widehat{N}_1, \widehat{N}_2, \widehat{H}) = ((2b - d)/K, 0, 0), (0, (2b - d)/K, 0))$ occurs in blue areas. We then compute the equilibria where coexistence of two species is possible when resource competition between species (e–h) cannot occur ($\alpha = 0$) and occurs (i–l) ($\alpha = 0.6$). Black, white, red, and blue areas represent alternative stable states, all-coexistence, competitive exclusion of a rarer species, and frequency-dependent coexistence. Parameters are $b = 1$, $d = d_h = 0.1$, $x_1 = 0$, $x_2 = 2$, $x_{\mathrm{h}} = 1$, and $K = 1000$.

### Supplementary figure 4

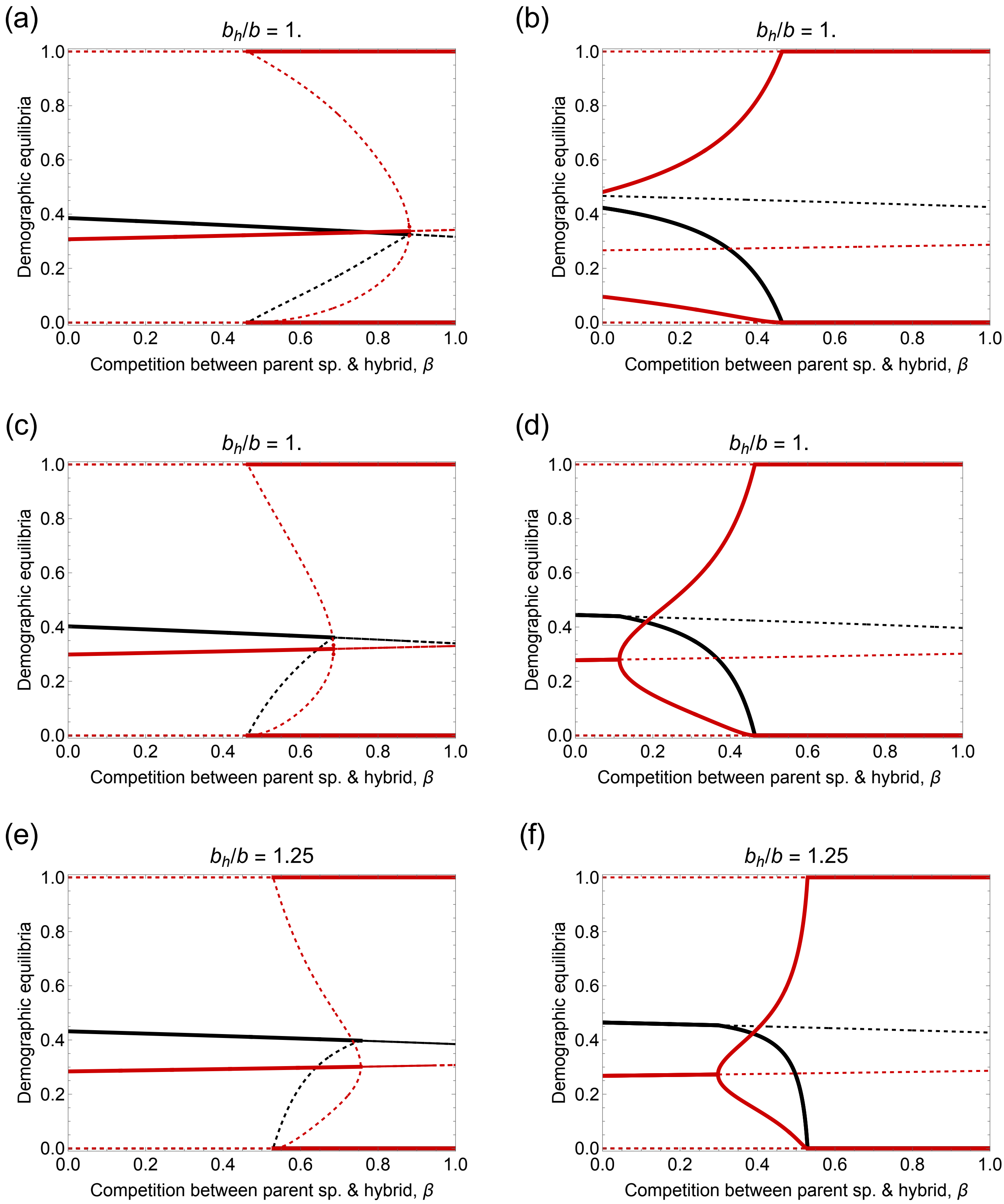


**Figure S4**: Bifurcation diagrams of demographic equilibria of resident species 1 (red) and hybrid (black). We show the frequencies of resident species and hybrid at equilibrium along the change in the strength of competition between parent species and hybrid. Solid and dashed lines denote locally stable and unstable equilibria, respectively. Parameters of panels (a–f) are corresponding to the panels (b, c, e, f, h, i) in Fig. 1, respectively.

### Supplementary figure 5

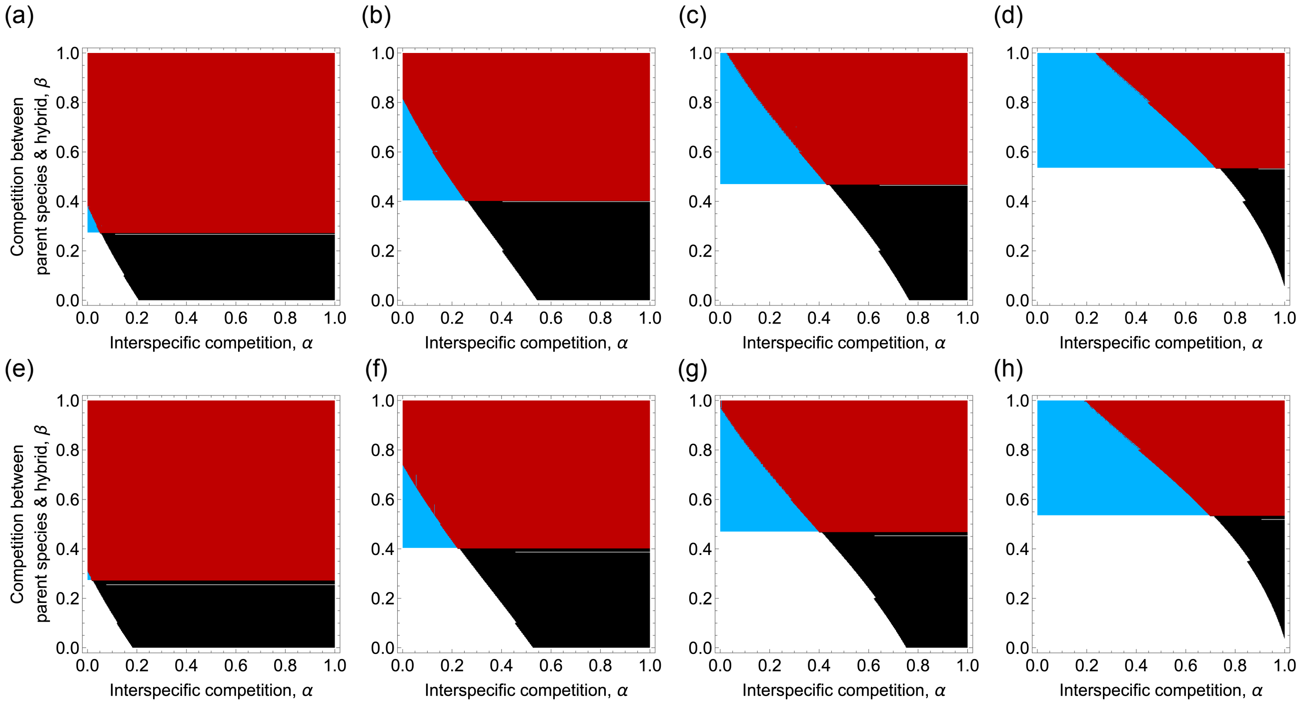


**Figure S5**: The dependence of ecological outcomes on the relative birth rate of hybrid versus parent species. Panels (a–c) assume no search cost ($e = 0$); panels (d–f) include a search cost ($e = 450$). (a–d) When the hybrid birth rate is close to that of parent species, the four outcomes described in the main text emerge. As the hybrid birth rate decreases, the parameter space for frequency-dependent coexistence (blue area) narrows. (e–h) In the presence of the searching cost, we obtained qualitatively the same results. Black, white, red, and blue areas represent alternative stable states, all-coexistence, competitive exclusion of a rarer species, and frequency-dependent coexistence. Parameters are $b_{\mathrm{h}} =$ (a, e) 0.25, (b, f) 0.75, (c, g) 1, and (d, h) 1.25. Other parameters are $b = 1$, $d = d_{\mathrm{h}} = 0.1$, $\sigma_r^2 = 25$, $x_1 = 0$, $x_2 = 2$, $x_{\mathrm{h}} = 1$, and $K = 1000$.

### Supplementary figure 6

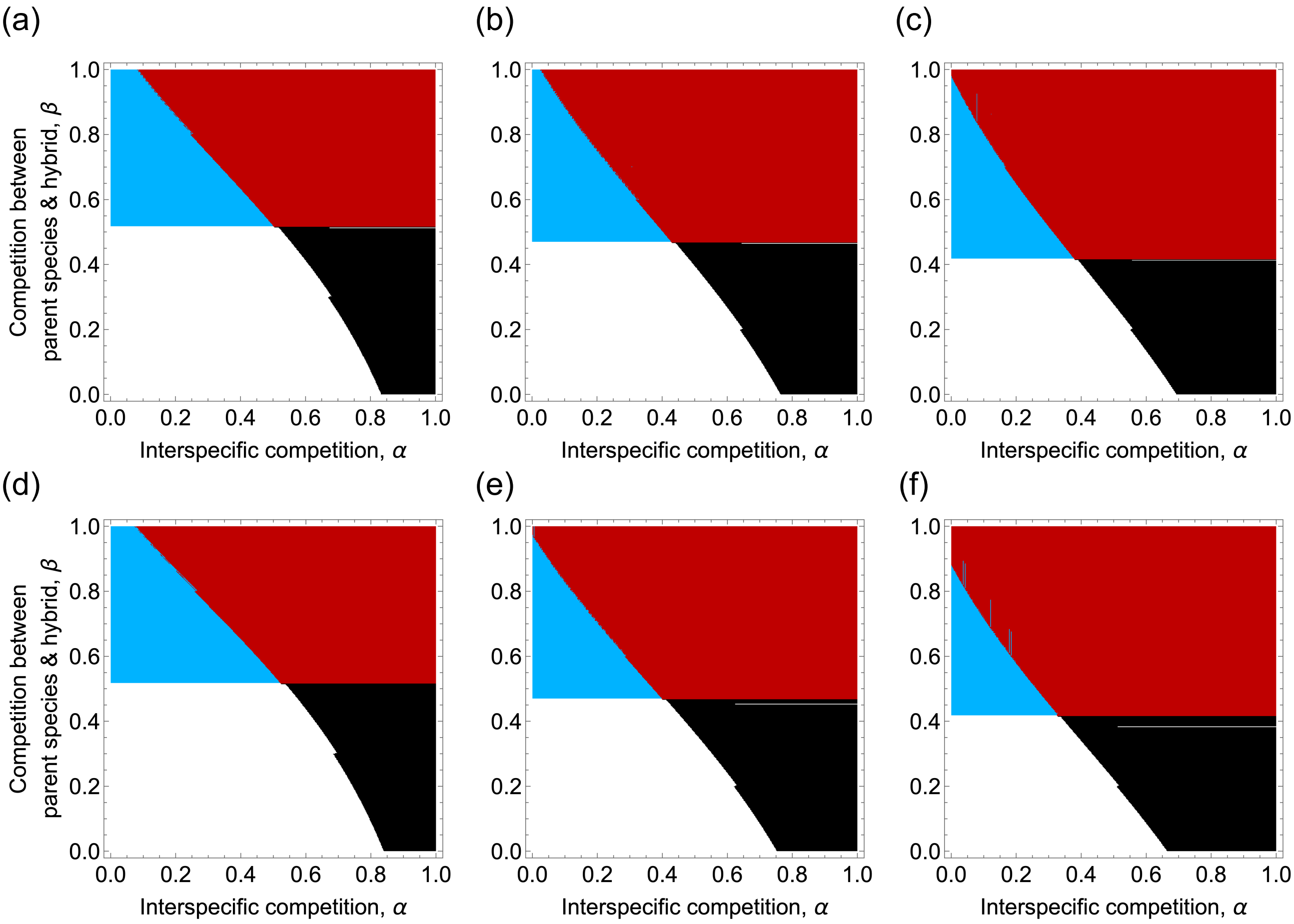


**Figure S6**: The dependence of ecological outcomes on the relative mortality rate of hybrid versus parent species. The outcomes are qualitatively the same as those when we change the relative birth rate of hybrid over parent species. Panels (a–c) assume no search cost ($e = 0$); panels (d–f) include a search cost ($e = 450$). (a–c) Higher hybrid mortality expands the parameter space leading to extinction of the rare species (blue and red areas). (d–f) In the presence of the searching cost, we obtained qualitatively the same results. Black, white, red, and blue areas represent alternative stable states, all-coexistence, competitive exclusion of a rarer species, and frequency-dependent coexistence. Parameters are $d_{\mathrm{h}} =$ (a, d) $0.01$, (b, e) $0.1$, and (c, f) $0.2$. Other parameters are $b = b_{\mathrm{h}} = 1$, $d = 0.1$, $\sigma_r^2 = 25$, $x_1 = 0$, $x_2 = 2$, $x_{\mathrm{h}} = 1$, and $K = 1000$.

### Supplementary figure 7

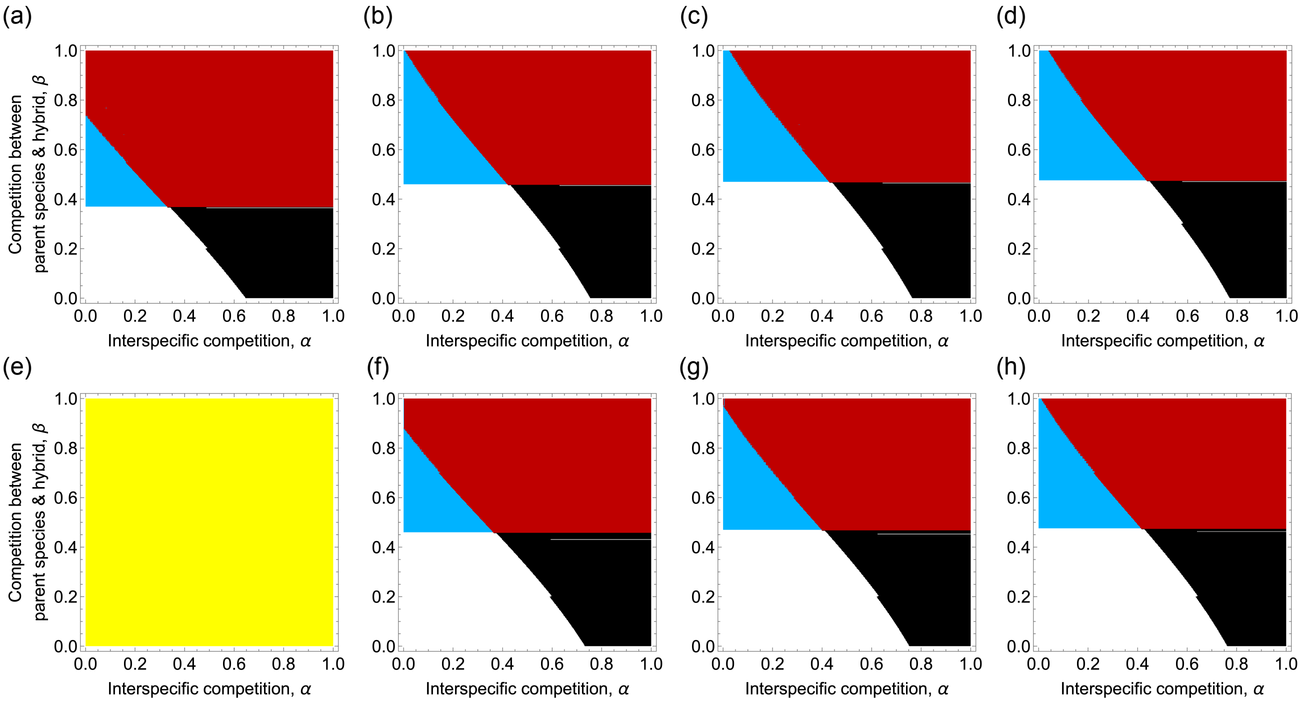


**Figure S7**: The dependence of ecological outcomes on equal birth rates of parent species and hybrids. Panels (a–c) assume no search cost ($e = 0$); panels (d–f) include a search cost ($e = 450$). (b–d, f–h) High birth rates yield qualitatively similar results. (a) When birth rates are low, extinction of the rare species is more likely. Specifically, (e) both species and their hybrid cannot persist when searching for mating partners is costly. Black, white, red, blue, and yellow areas represent alternative stable states, all-coexistence, competitive exclusion of a rarer species, frequency-dependent coexistence, and all-extinction. Parameters are $b = b_\mathrm{h} =$ (a, e) $0.25$, (b, f) $0.75$, (c, g) $1$, and (d, h) $1.25$. Other parameters are $d = d_\mathrm{h} = 0.1$, $\sigma_r^2 = 25$, $x_1 = 0$, $x_2 = 2$, $x_\mathrm{h} = (x_1 + x_2)/2$, and $K = 1000$.

Supplementary figure 8

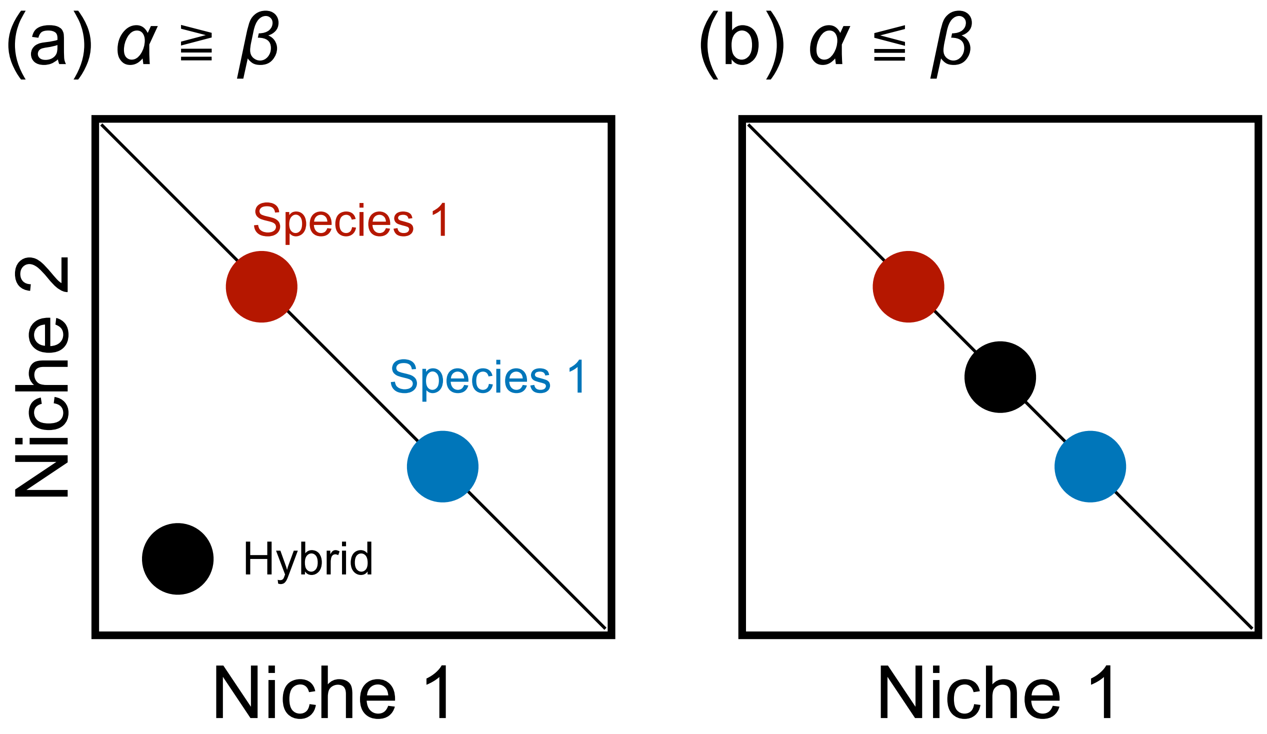


**Figure S8**: the relative strength of interspecific competition to competition between parent species and their hybrid is determined by the combination of niche traits. (a) When hybrid obtained distant niche from those of parent species through hybridization, interspecific competition is more severe than hybrid-parent competition. (b) When hybrid acquired intermediate niche between those of parent species, interspecific competition is weaker than hybrid-parent competition.

### Supplementary table

Table S1: Probability of genotype reproduction from hybrid×hybrid mating.

| Parental haplotype | A | a |
|---|---|---|
| A | AA | Aa |
| a | Aa | aa |
